# Agricultural fertilization with poultry manure results in persistent environmental contamination with the pathogen *Clostridioides difficile*

**DOI:** 10.1101/2021.02.01.429155

**Authors:** Martinique Frentrup, Nadine Thiel, Vera Junker, Wiebke Behrens, Steffen Münch, Paul Siller, Tina Kabelitz, Matthias Faust, Alexander Indra, Stefanie Baumgartner, Kerstin Schepanski, Thomas Amon, Uwe Roesler, Roger Funk, Ulrich Nübel

## Abstract

During a field experiment applying broiler manure for fertilization of agricultural land, we detected viable *Clostridioides* (formerly, *Clostridium*) *difficile* in broiler feces, manure, dust, and fertilized soil. A large diversity of toxigenic *C. difficile* isolates was recovered, including PCR ribotypes common from human disease. Genomic relatedness of *C. difficile* isolates from dust and from soil, recovered more than two years after fertilization, traced their origins to the specific chicken farm that had delivered the manure. We present evidence of long-term contamination of agricultural soil with manure-derived *C. difficile* and demonstrate the potential for airborne dispersal of *C. difficile* through dust emissions during manure application. *Clostridioides* genome sequences virtually identical to those from manure had been recovered from chicken meat and from human infections in previous studies, suggesting broiler-associated *C. difficile* are capable of zoonotic transmission.

## Introduction

The anaerobic gut bacterium *Clostridioides difficile* (formerly, *Clostridium difficile* (Lawson *et al*., 2016)) is the most frequent infectious cause of antibiotic-associated diarrhea and among the leading culprits of healthcare-associated infections (Martin *et al*., 2016). However, modelling studies have suggested that transmission in the community and in the healthcare system were equally relevant for sustaining *C. difficile* in the human population (Durham *et al*., 2016; McLure *et al*., 2019). Patients asymptomatically colonized with *C. difficile* upon hospital admission have a six-fold increased risk of suffering a *C. difficile* infection (CDI) (Zacharioudakis *et al*., 2015), and even without developing CDI themselves they may increase the overall burden of nosocomial CDI significantly by spreading the pathogen to other patients (Longtin *et al*., 2016; Blixt *et al*., 2017; Donskey *et al*., 2018). In addition, CDI occurs independent from healthcare at increasing incidence (Ofori *et al*., 2018), but reservoirs and pathways of transmission outside of the hospital environment are incompletely understood (Warriner *et al*., 2017; Rodriguez Diaz *et al*., 2018).

Toxigenic *C. difficile* seems widespread in various environments, since it was recovered from domestic wastewater (Moradigaravand *et al*., 2018; Numberger *et al*., 2019) and river sediments (Zidaric *et al*., 2010), from retail compost (Lim *et al*., 2020), soil (Janezic *et al*., 2016) and root vegetables (Lim *et al*., 2018; Tkalec *et al*., 2019). It was also found to colonize various mammals and birds, including wildlife, pets, and livestock (Weese, 2020). Notably, fattening pigs have been proposed as a potential source for transmission of *C. difficile* to humans, since strains with highly related genomes were isolated from both, pigs and farm workers (Knetsch *et al*., 2018). *Clostridioides difficile* was also detected in chicken feces and chicken meat repeatedly (Zidaric *et al*., 2008; Weese *et al*., 2010; Harvey *et al*., 2011; Abdel-Glil *et al*., 2018; Heise *et al*., 2021), even though there is no evidence for significant CDI in birds (Weese, 2020). Livestock manure often contains *C. difficile* even after being treated by composting or fermentation in biogas plants (Usui *et al*., 2017; Dharmasena and Jiang, 2018; Le Maréchal *et al*., 2020). As a consequence, the disposal of manure or manure-derived products as fertilizer on agricultural land may lead to environmental contamination with *C. difficile* spores. The survival of *C. difficile* in fertilized agricultural soil and its release with surface water runoff or dust has as yet not been investigated, in contrast to other manure-derived pathogens (Blaustein *et al*., 2015; Thiel *et al*., 2020).

The spread of pathogenic bacteria can be tracked by comparing their genome sequences (Croucher *et al*., 2015; Besser *et al*., 2019; Thiel *et al*., 2020). Within the EnteroBase platform, we have recently established a publicly accessible database for *Clostridioides* genomic data that currently (January 2021) contains 20,972 draft genomes and their associated metadata (Frentrup *et al*., 2020). Standardized sequence data assembly and quality control in conjunction with core-genome multilocus sequence typing (cgMLST) and hierarchical clustering of cgMLST allelic profiles - as implemented in EnteroBase - facilitates the detection of *C. difficile* spread, since isolates from transmission chains frequently can be identified by being related at the HC2 level (i.e. constituting chains of genomes with pairwise differences of maximally two cgMLST alleles) (Frentrup *et al*., 2020). Moreover, widespread epidemic strains commonly are related at the HC10 level, and PCR ribotypes correlate well with clusters at the HC150 level (which we dubbed ‘core-genome sequence typing complexes’; CC) (Frentrup *et al*., 2020).

In the present study, we detected the persistence of viable *C. difficile* in agricultural soil for several years following its fertilization with manure from broiler chickens. Genomic relatedness of *C. difficile* isolates from soil and from dust released during the fertilization process traced their origins to the specific chicken farm that had delivered the manure.

## Results

### Diversity of C. difficile isolates in chicken manure

Chicken manure was sampled at three different locations, including two farms and a manure trading cooperative. Altogether 146 *C. difficile* isolates were obtained from manure samples by applying an anaerobic enrichment protocol (Janezic *et al*., 2018) and their genomes were sequenced. Genomic data indicated that 98% of the isolates carried both toxin genes, *tcdA* and *tcdB* (Suppl.Figure S1A), and only three isolates were non-toxigenic. Analysis of genome sequences with EnteroBase showed that manure isolates were related to 13 CCs (i.e. hierarchical clusters at the level HC150), which we had previously shown to correlate well with PCR ribotypes (RT, Figure 1A) (Frentrup *et al*., 2020). The majority of isolates (94%) from Farm 1 were related to CC3 (Table 1, Figure 1B), which corresponds to PCR ribotype 001 (Frentrup *et al*., 2020), and repeated samplings showed that this predominance of CC3 at Farm 1 was evident over a period of at least one year (Suppl. Figure S1B). In contrast, only one isolate (4%) from Farm 2 was CC3, and none from the manure trader (Figure 1B). Instead, isolates from the latter two suppliers were distributed among a number of different CCs, the most predominant of which were CC71 (RT014/020), CC88 (RT014), CC2 (RT002), CC86 (RT005), and CC391 (RT081) (Figure 1).

**Table 1.** Core-genome sequence type complexes (CC)

| **CC** | **PCR Ribotype** | **Farm1** | **Farm2** | **trader** |
| --- | --- | --- | --- | --- |
| 2 | 002 | 4 (4.3 %) | 1 (4.2 %) | 7 (25 %) |
| 3 | 001 | 88 (93.6 %) | 1 (4.2 %) | 0 |
| 71 | 014/020 | 0 | 10 (41.7 %) | 3 (10.7 %) |
| 86 | 005 | 1 (1.1 %) | 2 (8.3 %) | 4 (14.3 %) |
| 88 | 014 | 0 | 2 (8.3 %) | 10 (35.7 %) |
| 391 | 081 | 0 | 5 (20.8 %) | 0 |
| 5408 | 029 | 0 | 1 (4.2 %) | 0 |
| 5410 | novel | 0 | 2 (8.3 %) | 0 |
| 207 | 003 | 1 (1.1 %) | 0 | 0 |
| 34 | 014 | 0 | 0 | 1 (3.6 %) |
| 596 | 011/049 | 0 | 0 | 1 (3.6 %) |
| 645 | 029 | 0 | 0 | 1 (3.6 %) |
| 1643 | 011/049 | 0 | 0 | 1 (3.6 %) |

**Figure 1.**
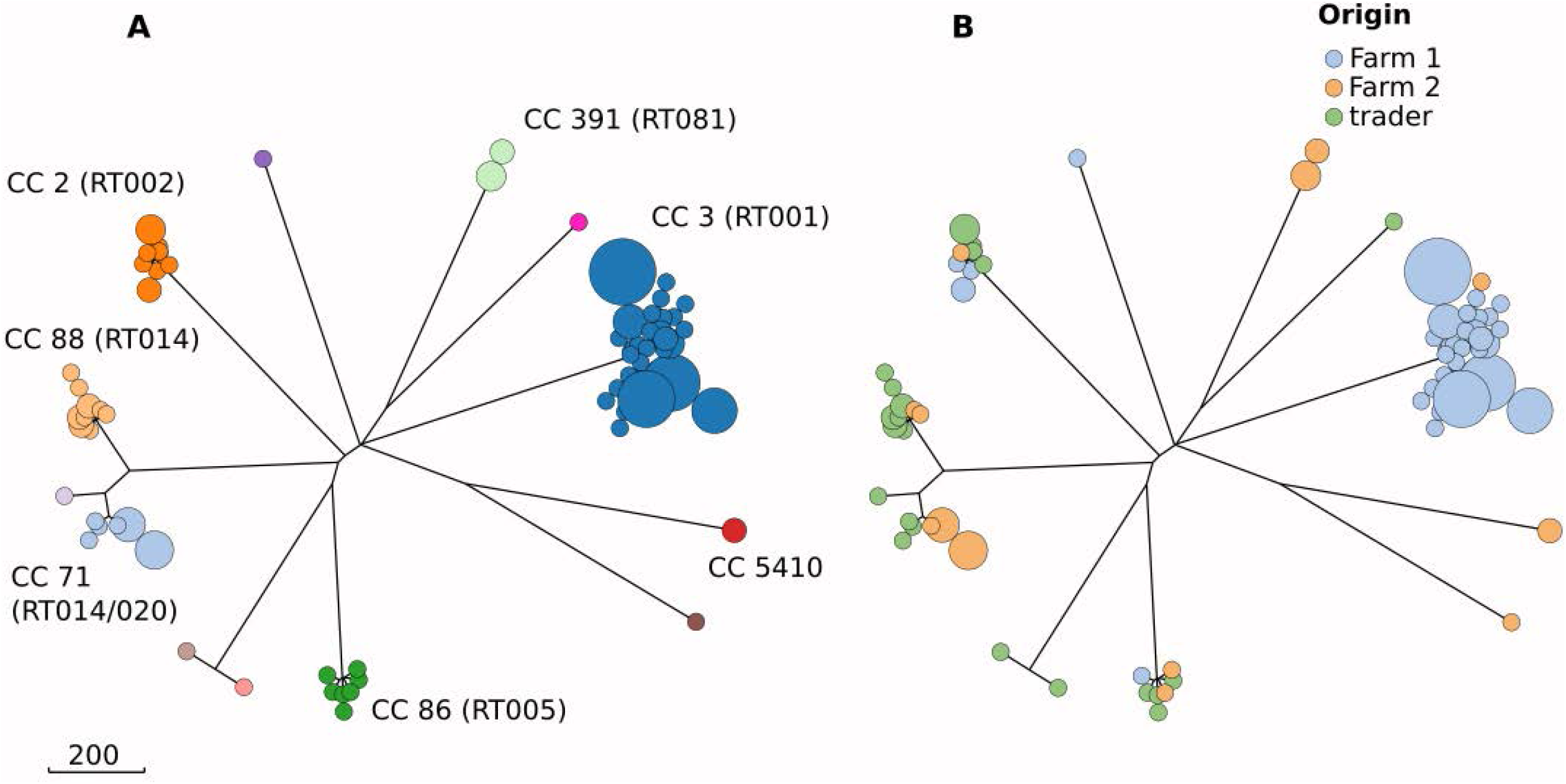
Rapid-neighbour-joining phylogenetic trees based on cgMLST allelic differences between *C. difficile* isolates (n=146) from manure samples. The scale bar indicates the branch length corresponding to sequence differences at 200 cgMLST loci. **A** Colors indicate CCs (core-genome sequence-typing complexes); RT, PCR ribotype. **B** Colors indicate origins of manure.

### Close genomic relationships identify source of environmental C. difficile

A 2.1-hectare agricultural field was fertilized with 12 tons of poultry manure from Farm 1 (Thiel *et al*., 2020). Prior to fertilization, our enrichment approach failed to detect any *C. difficile* in soil from this field. After fertilization, however, *C. difficile* got enriched and cultivated consistently from soil samples collected at multiple points in time for up to 143 weeks (Figure 2). Moreover, one dust sample collected during manure spread by using an aerosol collection device (Thiel *et al*., 2020) at the edge of the field tested positive for *C. difficile* by enrichment (Figure 2). Of note, *C. difficile* was detected in soil and dust by enrichment culture only, whereas cultivation and quantification by direct plating on selective agar medium was not successful. Altogether, we collected 144 *C. difficile* isolates from fertilized soil and from dust, and from poultry feces and manure from Farm 1. Bacterial genome sequencing and cgMLST-based hierarchical clustering analysis with EnteroBase (Frentrup *et al*., 2020) resulted in three HC2 clusters (HC2_1232, HC2_5435, HC2_5465; Figure 3) and four singletons. Generally, hierarchical clustering at the level HC2 indicates close genomic relationships of *C. difficile* isolates; it was previously shown to correlate with events of transmission between hospital patients (Frentrup *et al*., 2020).

**Figure 2.**
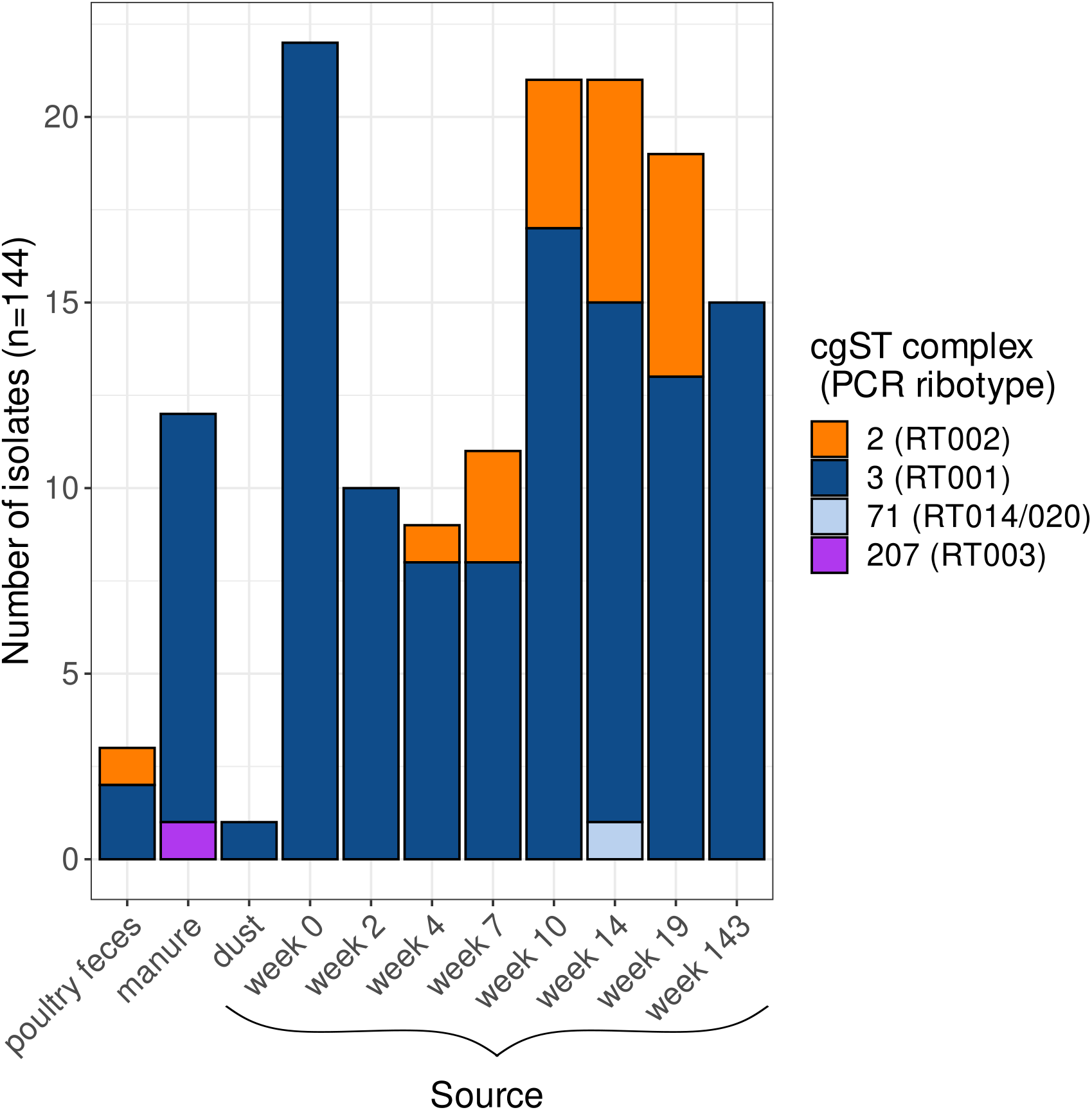
Distribution of isolates recovered from chicken feces and manure from Farm 1 and from samples collected during the field experiment (n=144). Colors represent CCs, which were determined based on cgMLST allelic profiles in EnteroBase.

**Figure 3.**
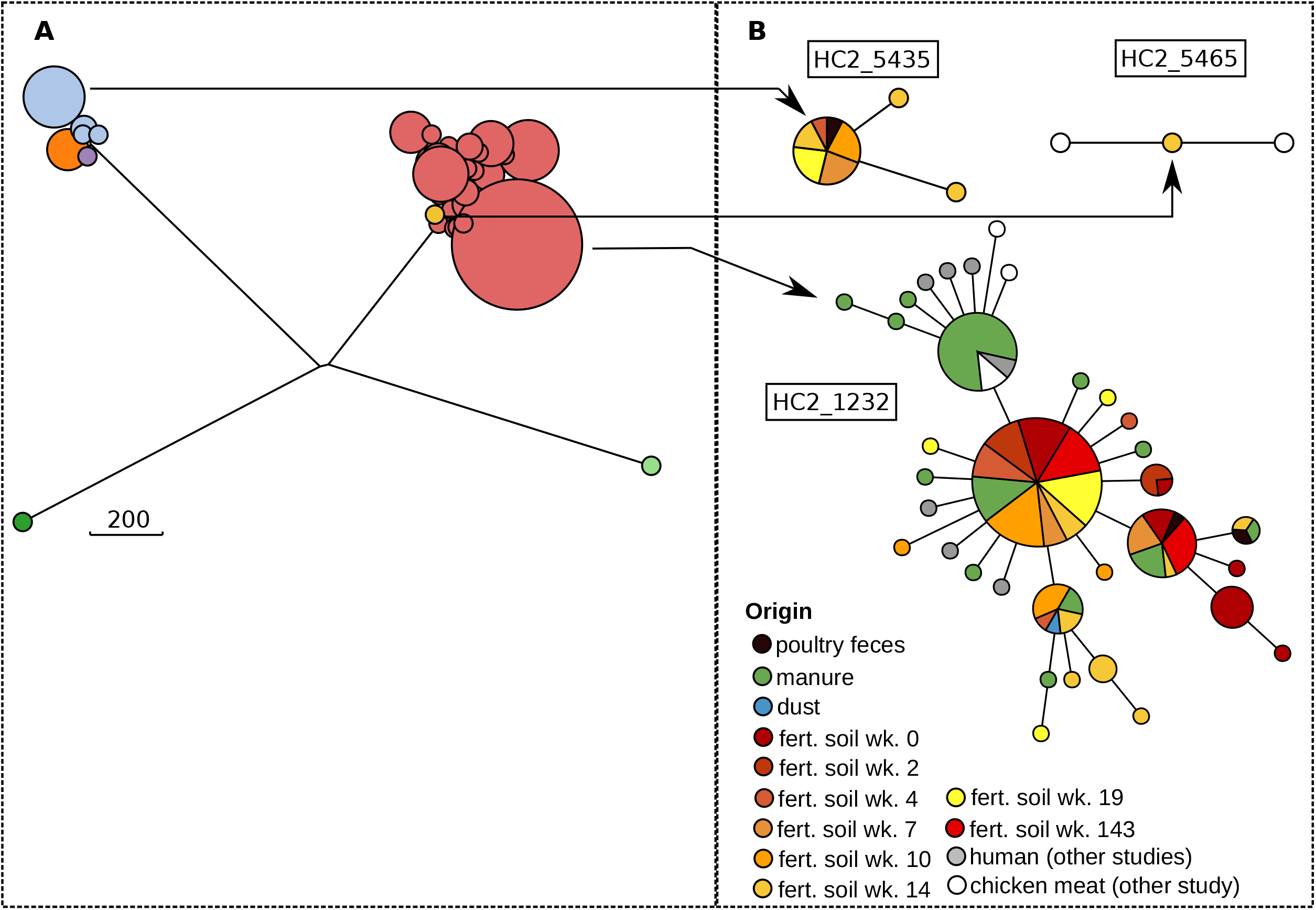
**A** Rapid-neighbor-joining phylogenetic tree based on cgMLST allelic profiles from all isolates (n=144) sampled during the field experiment. Colors indicate HC2 clusters. The scale bar indicates the branch length corresponding to sequence differences at 200 cgMLST loci. **B** Minimum-spanning trees for two HC2 clusters. Numbers on branches indicate the number of cgMLST allelic differences and the colors represent the source from which the isolates were extracted from.

Two HC2 clusters (HC2_1232, HC2_5435) included genomes from two or more different sources, including chicken feces collected at Farm 1, manure from Farm 1, dust collected during the application of manure to the field, and fertilized soil from multiple points in time (Table 3 and Figure 3). This result confirmed that the *C. difficile* strains that were recovered during and after fertilization indeed originated from Farm 1, i.e. they had been disseminated onto the agricultural field through the fertilization process. These close genomic relationships were found among *C. difficile* isolates from all soil samples, indicating the persistence of viable, manure-derived *C. difficile* in the soil for up to 143 weeks after fertilization (Figure 3). Likewise, the detection of closely related *C. difficile* in mineral dust showed that viable cells of the pathogen got aerosolized during the fertilization process and transported in an ascending dust plume at a distance of at least 20 meters from the applying tractor (Figure 3).

**Table 2.** Genotypes and antibiotic susceptibilities of 19 selected *C. difficile* isolates

|  |  |  |  | MIC** [µg/ml] |  |  |  |  | Gene content (predicted***\|PCR) |  |  |  |
| --- | --- | --- | --- | --- | --- | --- | --- | --- | --- | --- | --- | --- |
| Isolate | Source | Origin | CC (RT predicted*\|PCR) | VAN | MTZ | MXF | CLI | TET | *tcdA* | *tcdB* | *cdtA* | *cdtB* |
| CD-17-00892 | manure | trader | 88 (RT014\|RT014) | 0.5 | 0.094 | 0.75 | 1.5 | 0.016 | ++ | ++ | - | - |
| CD-17-01035 | manure | trader | 1643 (RT011/049\|RT049) | 0.5 | 0.125 | 0.75 | 3 | 0.047 | ++ | ++ | - | - |
| CD-17-01037 | manure | trader | 34 (RT014\|RT014) | 0.5 | 0.125 | 0.75 | 2 | 0.032 | ++ | ++ | - | - |
| CD-17-01039 | manure | trader | 645 (RT029\|RT029) | 0.75 | 0.125 | 0.75 | 0.5 | 0.047 | ++ | ++ | - | - |
| CD-17-01040 | manure | trader | 596 (RT011/049\|RT049) | 0.5 | 0.125 | 0.75 | 3 | 0.047 | ++ | ++ | - | - |
| CD-17-01068 | manure | Farm 1 | 207 (RT003\|RT003) | 0.38 | 0.094 | 1 | 0.5 | 0.023 | ++ | ++ | - | - |
| CD-17-01070 | dust | field experiment | 3 (RT001\|RT001) | 0.5 | 0.094 | 0.5 | 1.5 | 0.032 | ++ | ++ | - | - |
| CD-17-01381 | manure | Farm 2 | 71 (RT014/020\|RT014) | 0.75 | 0.125 | 0.75 | 1.5 | 0.75 | ++ | ++ | - | - |
| CD-17-01390 | manure | Farm 2 | 391 (RT081\|RT081) | 0.5 | 0.125 | 0.75 | 2 | 0.032 | ++ | ++ | - | - |
| CD-17-01395 | manure | Farm 2 | 5408 (n.a.\|RT029) | 0.75 | 0.125 | 1 | 3 | 0.032 | - | - | - | - |
| CD-17-01424 | manure | Farm 2 | 5410 (n.a.\|novel) | 0.38 | 0.032 | 1 | 2 | 0.023 | - | - | - | - |
| CD-17-01524 | manure | Farm 1 | 86 (RT005\|RT005) | 1 | 0.19 | 0.75 | 6 | 0.032 | ++ | ++ | - | - |
| CD-18-00685 | manure | Farm 1 | 2 (RT002\|RT002) | 0.5 | 0.064 | 1 | 2 | 0.023 | ++ | ++ | - | - |
| CD-19-00355 | fert. soil wk. 7 | field experiment | 2 (RT002\|RT002) | 0.5 | 0.125 | 0.75 | 2 | 0.064 | ++ | ++ | - | - |
| CD-19-00409 | fert. soil wk. 14 | field experiment | 71 (RT014/020\|RT014) | 0.75 | 0.125 | 1 | 4 | 0.032 | ++ | ++ | - | - |
| CD-19-00417 | fert. soil wk. 14 | field experiment | 3 (RT001\|RT001) | 0.5 | 0.19 | 0.5 | 1.5 | 0.047 | ++ | ++ | - | - |
| CD-19-00426 | fert. soil wk. 19 | field experiment | 2 (RT002\|RT002) | 0.38 | 0.25 | 1 | 4 | 0.047 | ++ | ++ | - | - |
| CD-19-00513 | manure | Farm 1 | 3 (RT001\|RT001) | 0.75 | 0.75 | 0.5 | 4 | 0.047 | ++ | ++ | - | - |
| CD-20-00542 | fert. soil wk. 143 | field experiment | 3 (RT001\|RT001) | 0.75 | 0.75 | 0.75 | 6 | 0.047 | ++ | ++ | - | - |

\* Ribotypes were predicted based on hierarchical clustering in EnteroBase
\*\* VAN: vancomycin; MTZ: metronidazole; MXF: moxifloxacin; CLI: clindamycin; TET: tetracycline; *tcdA*: gene encoding for toxin A; *tcdB*: gene encoding for toxin B; *cdtA* and *cdtB*: genes encoding for the binary toxin
\*\*\* Toxin gene prediction was based on corresponding wgMLST loci (+=present; -=absent)

**Table 3.** HC2 clusters

| Source | HC2_1232 | HC2_4410 | HC2_5435 | HC2_5465 | HC2_12193 | HC2_12207 | HC2_12213 |
| --- | --- | --- | --- | --- | --- | --- | --- |
| poultry feces | 2 | 0 | 1 | 0 | 0 | 0 | 0 |
| manure | 11 | 1 | 0 | 0 | 0 | 0 | 0 |
| dust | 1 | 0 | 0 | 0 | 0 | 0 | 0 |
| fert. soil wk. 0 | 22 | 0 | 0 | 0 | 0 | 0 | 0 |
| fert. soil wk. 2 | 10 | 0 | 0 | 0 | 0 | 0 | 0 |
| fert. soil wk. 4 | 8 | 0 | 1 | 0 | 0 | 0 | 0 |
| fert. soil wk. 7 | 8 | 0 | 3 | 0 | 0 | 0 | 0 |
| fert. soil wk. 10 | 17 | 0 | 3 | 0 | 1 | 0 | 0 |
| fert. soil wk. 14 | 13 | 0 | 4 | 1 | 2 | 0 | 1 |
| fert. soil wk. 19 | 13 | 0 | 3 | 0 | 2 | 1 | 0 |
| fert. soil wk. 143 | 15 | 0 | 0 | 0 | 0 | 0 | 0 |

### PCR ribotypes and antibiotic susceptibilities

*Clostridioides difficile* isolates (n=19) selected to represent sources (i.e. manure from the different suppliers, fertilized soil, and dust) and genomic diversity (at the level of CCs) proved to be phenotypically susceptible to the antibiotics vancomycin, metronidazole, moxifloxacin, clindamycin, and tetracycline (Table 2). None of the genome sequences (n=278) carried resistance-causing mutations in the gyrase gene *gyrA* (Zaiß *et al*., 2010), confirming the lack of fluoroquinolone resistance in our strain collective (not shown). PCR ribotypes determined in the laboratory were fully concordant with ribotype predictions based on hierarchical clustering in EnteroBase (Table 2).

### Closely related clinical and poultry meat isolates

Hierarchical clustering of cgMLST allelic profiles in EnteroBase routinely determines genomic relationships at multiple phylogenetic levels among all >20,000 entries in the *Clostridioides* database (Frentrup *et al*., 2020). Remarkably, a limited number of genome sequences from several previous studies were closely related (at HC2 level) to those from Farm 1 (Figure 3). Most notably, virtually identical *C. difficile* genome sequences had been recovered from retail chicken meat (n=6; Figure 3), which had been purchased in one region in Germany (Berlin and Brandenburg), but had been produced in a number of different cutting plants in Germany and the Netherlands (Heise et al., 2021). Additional closely related genomes originated from isolates from human patients suffering from C. difficile infection in Germany (n=2), the Netherlands (n=5) and Hungary (n=1; Figure 3). Of note, these genomic similarities were not due to impaired quality of the sequence data, since >99% of cgMLST alleles were successfully called for all genome sequences. Moreover, no genes of the whole-genome MLST set (Frentrup *et al*., 2020) were differentially present (not shown), indicating that accessory genomes were virtually identical among all these isolates, too,

## Discussion

### Chicken manure carried diverse C. difficile, including clinically relevant strains

Almost all *C. difificile* isolates from manure in our study carried the *tcdA* and *tcdB* genes in their genomes, and hence must be considered fully virulent and able to cause gastrointestinal disease in humans. This result is in concordance with most previous studies on poultry-associated *C. difficile* (e.g. (Dharmasena and Jiang, 2018; Berger *et al*., 2020; Le Maréchal *et al*., 2020; Heise *et al*., 2021)) even though there is little evidence that *C. difficile* may cause disease in birds (Weese, 2020).

In manure samples from three suppliers, we found a total of 13 CCs (core-genome sequence-type complexes) of *C. difficile*. CCs correlate well with PCR ribotypes (Frentrup *et al*., 2020) (Table 2), and ribotypes 001, 014/020 and 005 have been reported from poultry feces (Indra *et al*., 2009; Hussain *et al*., 2016; Abdel-Glil *et al*., 2018; Le Maréchal *et al*., 2020) and from broiler meat (De Boer *et al*., 2011; Tkalec *et al*., 2020) in the past. In our manure samples, the most predominant strains were CC3 (RT001), CC71 (RT014/020) and CC88 (RT014). Remarkably, these are also among the most prevalent strains causing human *C. difficile* infections in Europe (Davies *et al*., 2016). However, our isolates from broiler chickens were not resistant to fluoroquinolones or clindamycin, in contrast to the vast majority of clinical RT001 isolates from human CDI (Zaiß *et al*., 2010; Eyre *et al*., 2018). This striking difference in antibiotic resistances suggests that *C. difficile* RT001 in chickens constitutes a population separate from the epidemic RT001 strain causing healthcare-associated CDI in humans, with limited exchange. This notion was confirmed by hierarchical clustering, which indicated that all our CC3 *C. difficile* from broiler manure (n=199) were related to a single HC10 cluster (HC10_783; Suppl. Table 1) that currently includes only 15 (7%) human-associated *C. difficile* isolates in EnteroBase. This separation was not observed for RT014/020, which is antibiotic resistant more rarely (Zaiß *et al*., 2010; Eyre *et al*., 2018), and where 13 isolates from broilers were affiliated to nine different HC10 clusters (Suppl. Table 1), the larger of which included numerous isolates from diverse host species and geographic origins. Fluoroquinolone and clindamycin resistance in poultry-associated *C. difficile* has occasionally been reported (from the USA and Zimbabwe (Harvey *et al*., 2011; Dharmasena and Jiang, 2018; Berger *et al*., 2020)). Since macrolides and fluoroquinolones are the two antibiotics most heavily used in the poultry industry in Europe, and resistance against these drugs is widespread among other gastrointestinal pathogens from chickens (Roth *et al*., 2019), lowered susceptibilities might also have been expected from broiler-associated *C. difficile*, but yet this was not detected in our samples.

### Long-term persistence of manure-derived C. difficile in fertilized agricultural soil

*Clostridioides difficile* has been reported from a wide range of different environmental samples, including soil (Rodriguez Diaz *et al*., 2018). To our best knowledge, however, our study is the first to use genome sequence analysis to trace environmental *C. difficile* back to its source. As one result, we show that *C. difficile i*n fertilized soil indeed originated from chickens in Farm 1. Hence, our field experiment demonstrated that manure-derived *C. difficile* remained viable in fertilized soil over the entire study period, i.e. for at least 143 weeks, or almost three years. The continued bacteriological detection of *C. difficile* in all samples investigated suggested that its survival may be much longer than the sampling period, even though precise extrapolation was not possible due to the failure of quantitative cultivation. The observed long-term contamination of the soil certainly was enabled by the ability of *C. difficile* to produce endospores, which can stay viable for many years (Yang and Ponce, 2011). In contrast, these bacteria are unlikely to perform much metabolic activity or even proliferate under ambient conditions in the soil, since their physiology is adapted to life in the intestines of warm-blooded animals.

We previously reported that chicken manure carried additional pathogens, including *Enterococcus faecium* and extended-spectrum beta-lactamase (ESBL) producing *Escherichia coli* (Thiel *et al*., 2020). However, ESBL *E. coli* died off within a few days during manure storage (Siller *et al*., 2020) and enterococci rapidly declined in soil within weeks after fertilization (Thiel *et al*., 2020). In the present study, in contrast, we demonstrate that viable *C. difficile* remained detectable in fertilized soil for several years and hence represented a long-lasting pollution.

### Potential for long-distance dispersal of C. difficile

Hierarchical clustering indicated that altogether 13 entries in the *Clostridioides* database shared identical HC2 clusters (HC2_1232, HC2_5465) with isolates from Farm 1, i.e. they had highly similar cgMLST profiles with at most two allelic differences, despite their origins from unrelated, previous studies. Seven of these isolates had been recovered from retail chicken meat from various cutting plants in Germany and the Netherlands (Heise *et al*., 2021), indicating widespread dissemination of *C. difficile* HC2_1232 by the poultry industry. Furthermore, the occurrence of the same HC2 clone in human CDI in Germany, Hungary and the Netherlands indicates that this strain is able to cause human disease. Consequently, this *C. difficile* HC2 clone poses a risk of zoonotic transmission.

It should be noted that pathogen genomic similarity alone does not prove direct transmission between remote places, but should be interpreted with particular care in the absence of additional, epidemiological evidence (Besser *et al*., 2019). However, several plausible scenarios for long-distance transport of poultry-associated *C. difficile* exist. Chicken meat contaminated with *C. difficile* (De Boer *et al*., 2011; Harvey *et al*., 2011; Candel-Pérez *et al*., 2020; Heise *et al*., 2021) gets distributed to customers through widely ramified retail chains. Similarly, pork products (i.e., meat or manure) were suspected to promote the long-distance spread of *C. difficile*, after closely related *C. difficile* genomes had been detected in fattening pigs and humans across large geographic distances, without any documented epidemiological connections (Knetsch *et al*., 2018; Knight *et al*., 2019). Another potential path for the long-range dissemination of livestock-associated *C. difficile* may be the transport of colonized, live animals, e.g. from farms to slaughterhouses (Heise *et al*., 2021). Potentially even more important is the globalized structure of the poultry industry, which ships industrially produced broiler chicks by airfreight for stocking fattening farms globally (Lowder *et al*., 2009). It would be interesting to investigate the colonization status of chickens upon their arrival at fattening farms.

In addition, here we show that mineral dust from agricultural operations may carry aerosolized, manure-derived *C. difficile*. This dust may stay airborne for several days and during this time may get transported over several hundred kilometers, depending on atmospheric conditions (Faust *et al*., 2020; Thiel *et al*., 2020). Poultry manure is particularly prone to aerosolization due to its high dry-matter content (Kabelitz *et al*., 2020; Thiel *et al*., 2020) and therefore, its application for fertilization of agricultural fields likely contributes to the airborne dispersal of chicken-associated *C. difficile* over long distances. Aerosolized *C. difficile* is considered a potential source of human infection when inhaled (Best *et al*., 2010), similar to other enteric pathogens (Jahne *et al*., 2015). Hence, *C. difficile* in agricultural dust may represent a risk of airborne zoonotic transmission. Taken together, our results corroborate the relevance of a ‘One Health’ approach for curbing the spread of *C. difficile* between human, livestock, and environmental reservoirs.

## Experimental procedures

### Manure samples

To capture the diversity of *C. difficile* isolates in manure samples, three samplings were performed on three different sites. Manure samples from two broiler fattening farms and one manure trading cooperative were investigated. In addition, chicken feces were sampled by collecting 30 chicken droppings from each of 11 stables in Farm 1. Farm 1 is an intensive poultry-fattening farm in Brandenburg, Germany, housing about 19,000 animals per stable on wood pellets. Manure from this farm was sampled three times (May 30th 2017, November 8th 2017 and May 19th 2018). In Farm 2, which is located in Saxony-Anhalt, Germany, manure was collected in four different stables on August 14th 2017. Manure from the trader was sampled on March 27th 2017.

### Field experiment

In a field experiment, 12 tons of chicken manure from Farm 1 (see above) were applied to a 2.1-hectare agricultural field, which had not been fertilized with animal manure for 15 years. Details of this experiment have been published previously (Thiel *et al*., 2020). Briefly, dust particles that were released during the fertilization process were collected by impingement into 5 mL phosphate-buffered saline (PBS) at a height above ground of 1.50 m and at a distance from the tractor of 20, 50, and 100 m, respectively. Soil samples were taken on three representative sites on the field site prior to fertilization, directly after, and two, four, seven, ten, 14, 19 and 143 weeks later.

### Isolation of C. difficile isolates

Ten g of poultry feces, manure and soil samples were mixed with 90 g Luria-Bertani broth (Roth) each and subsequently homogenized for 30 s with a bag mixer (Interscience). After sedimentation of coarse particles (30 min, room temperature), supernatants and impingement suspensions from the aerosol collector were diluted to extinction with PBS and subsequently streaked on ChromID *C. difficile*-agar (Biomérieux). After incubation at 37°C for 24 h, *C. difficile* colonies were identified by species-specific PCR (locus TR10) (Zaiß *et al*., 2009). In addition, for enrichment cultures, 0.5 mL of suspensions were added to 10 mL brain heart infusion (BHI) broth (Roth) supplemented with 0.1% taurocholic acid (Sigma), 0.1% cysteine (Sigma) and *C. difficile* selective Supplement (Oxoid) in Hungate tubes (Janezic *et al*., 2018). After seven days of incubation at 37°C, an ethanol shock was performed by adding an equal amount of absolute ethanol to 0.5 mL culture and incubation for 1 h at room temperature. The culture was centrifuged at 2,500 x g for 5 minutes, the resulting cell pellet was resuspended in 200 µL PBS, and 100 µL were plated on ChromID *C. difficile*-agar and incubated at 37°C for 24 h. Again, bacterial colonies were tested by *C. difficile*-specific PCR (Zaiß *et al*., 2009).

### Antibiotic susceptibility testing

Isolates from agar-plates were transferred to anaerobic BHI broth (Roth) in Hungate tubes and grown for two days at 37°C. Subsequently, the culture was diluted 1:5 with PBS and 100 µl was spread on Columbia blood-agar (Oxoid). For each antimicrobial agent, an E-test strip was applied to the agar surface, followed by 24 hours of incubation at 37°C. The tests were interpreted visually by reading the minimum inhibitory concentration (MIC). MICs were determined for vancomycin, metronidazole, moxifloxacin (Biomérieux), clindamycin and tetracycline (Liofilchem). For interpretation, MIC breakpoints for antibiotic resistance were applied according to Pirš et al. (Pirš *et al*., 2013): metronidazole, ≥2 µg/mL; vancomycin, ≥2 µg/mL; moxifloxacin, ≥4 µg/mL; clindamycin, ≥8 µg/mL; tetracycline, ≥16 µg/mL.

### PCR ribotyping

PCR ribotyping of *C. difficile* isolates was performed as reported previously (Indra *et al*., 2008), applying capillary electrophoresis and the Webribo database (https://webribo.ages.at/).

### Whole genome sequencing

Genomic DNA was extracted by using the DNeasy Blood & Tissue kit (Qiagen), libraries were prepared as described previously (Steglich *et al*., 2018) and sequenced on an Illumina NextSeq 500 machine using a Mid-Output kit (Illumina) with 300 cycles. Illumina sequencing reads were uploaded to EnteroBase (http://enterobase.warwick.ac.uk/) and assembled with the embedded standardized pipeline (Frentrup *et al*., 2020). Thirty-two sequences did not pass the quality check in EnteroBase (Frentrup *et al*., 2020) and were excluded from further analyses. For 278 genomes, cgMLST allelic profiles (>99% complete) were determined and cgMLST-based hierarchical clustering performed using EnteroBase tools. To visualize genomic relatedness, rapid-neighbor-joining and minimum-spanning trees were calculated applying GrapeTree (Zhou *et al*., 2018; Frentrup *et al*., 2020). PCR ribotypes were predicted based on genomic relatedness at the level HC150 (i.e. hierarchical clusters of genome sequences with pairwise differences of maximally 150 cgMLST alleles; for details see (Frentrup *et al*., 2020)).

Sequences of the *gyrA* gene (cgMLST locus CD630_00060) were scanned for the mutations Thr-82-Ile and Asp-71-Glu, which are associated with fluoroquinolone resistance in *C. difficile* (Zaiß *et al*., 2010).

All genome sequencing data were submitted to the European Nucleotide Archive (ww.ebi.ac.uk/ena) under the study accession number PRJEB42049. A list of all analyzed genomes can be found in Supplementary Table1.

### Detection of toxin genes

DNA from selected isolates (n=19) was tested for the presence of toxin genes *tcdA, tcdB, cdtA* and *cdtB* by PCR (Persson *et al*., 2008). The presence or absence of toxin genes *tcdA* and *tcdB* was determined for all genomes in this study (n=275) based on allelic numbers for toxin gene loci in EnteroBase (i.e., allele number 0 was interpreted as absence of gene).

## Acknowledgements

We thank the farm owners for supporting this study, allowing sample collection at their facilities and donation of manure for experiments. We are grateful to the staff of the sequencing laboratory at DSMZ for excellent technical assistance and we appreciate support received from the Leibniz research alliance Infections ’21. This study was funded by the Leibniz Association (grant number: SAW-2017-DSMZ-2).

---

**Supplementay Figure S1.**
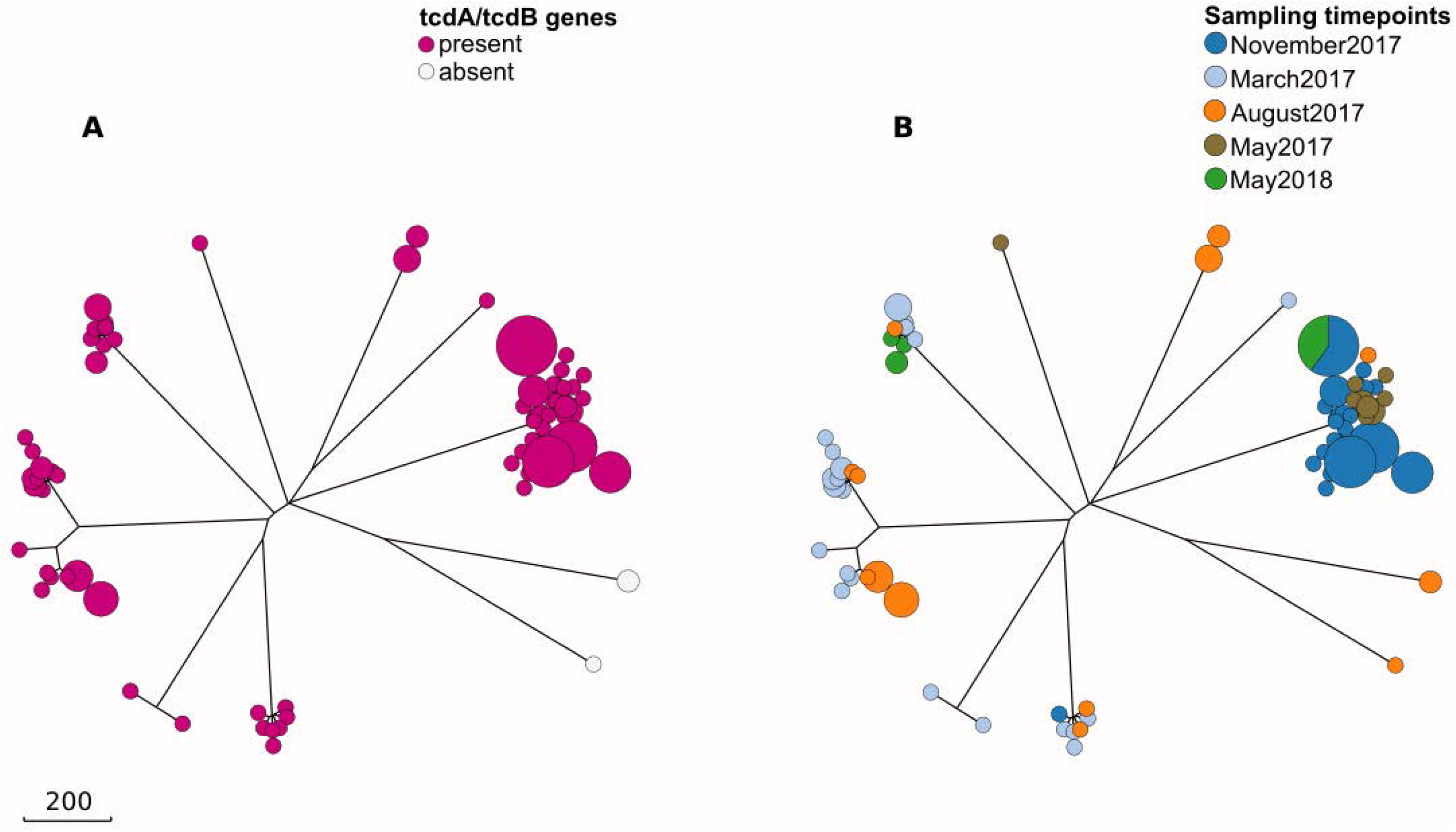
Rapid-neighbour-joining phylogenetic trees based on cgMLST allelic differences between *C. difficile* isolates (n=146) from manure samples (compare Figure 1). The scale bar indicates the branch length corresponding to sequence differences at 200 cgMLST loci. **A** Colors indicate the complement of toxin genes *tcdA* and *tcdB*. **B** Colors indicate sampling dates.

